# IL-22 Promotes Gammaherpesvirus Latency and Pathogenesis by Supporting Germinal Center Expansion and Polyclonal Autoimmunity

**DOI:** 10.64898/2026.02.20.706979

**Authors:** Sheikh Tahir Majeed, Samantha M. Bradford, Christopher N. Jondle

## Abstract

Gammaherpesviruses, including human Epstein-Barr Virus (EBV) and Kaposi’s Sarcoma-associated Herpesvirus (KSHV), are ubiquitous oncogenic pathogens linked to diverse malignancies, including Burkitt lymphoma, Hodgkin lymphoma, and nasopharyngeal carcinoma, as well as Kaposi’s sarcoma, as well as autoimmune disorders, most notably Multiple Sclerosis. These viruses colonize naïve B cells and drive a robust, polyclonal germinal center response to expand the latent viral reservoir and establish lifelong infection in memory B cells. Despite the clinical burden of these viruses, the host factors they usurp to maintain this chronic state remain poorly defined. Interleukin-22 (IL-22) is a critical cytokine traditionally recognized for its protective roles in bacterial and fungal defense; however, its involvement in gammaherpesvirus pathogenesis has not been previously explored. Using murine gammaherpesvirus 68 (MHV68) as a tractable *in vivo* model, we identify IL-22 as a novel proviral factor that promotes the establishment and maintenance of chronic infection. We show that MHV68 infection triggers a robust, T-cell-driven IL-22 response across primary and secondary sites of infection. Our studies in IL-22^-/-^ mice reveal a profound reduction in the virus-driven germinal center response, leading to diminished viral latency and impaired reactivation competency. Furthermore, we demonstrate that IL-22 supports the induction of the polyclonal, "irrelevant" antibody response, a signature of gammaherpesvirus-induced immunopathology. Together, our data reveal that gammaherpesviruses exploit the IL-22 axis to establish a supportive environment within the host, positioning this cytokine as a potential therapeutic target for disrupting the establishment of lifelong viral latency and its associated diseases.

**Author Summary:** Gammaherpesviruses are oncogenic double-stranded DNA viruses that establish lifelong infections in over 95% of adults worldwide and are multiple cancers, including many B-cell lymphomas. While the cytokine IL-22 is known for its role in clearing bacterial and fungal infections, its influence on chronic viral infections is poorly understood. In this study, we demonstrate for the first time that IL-22 plays a critical proviral role during gammaherpesvirus infection. By utilizing a murine model, we show that loss of IL-22 suppresses the establishment of viral latency, reduces viral reactivation and dampens the virus-driven germinal center response. viral latency and reactivation. Furthermore, we find that IL-22 signaling supports non-specific, polyclonal antibody surge characteristic of these infections. Our findings suggest that the virus exploits the IL-22 axis to shape a supportive microenvironment for its reservoir, identifying this pathway as a potential therapeutic target to limit viral latency and its associated immunopathology.

## Introduction

Gammaherpesviruses, including human Epstein-Barr virus (EBV) and Kaposi’s sarcoma-associated herpesvirus (KSHV), are ubiquitous pathogens that infect over 95% of the adult population worldwide and establish lifelong infection in the host (1-3). The gammaherpesviruses are etiological agents for a broad spectrum of malignancies, including cancers such as Burkitt’s lymphoma, Hodgkin’s lymphoma, and Kaposi’s sarcoma, particularly in immunocompromised individuals. Beyond oncology, chronic gammaherpesvirus infection has been increasingly linked to the development of systemic autoimmune diseases, most notably Multiple Sclerosis (4). The lifecycle of these viruses is characteristically biphasic, comprising an initial lytic replication phase (acute) followed by the establishment of viral latency (chronic phase) (5, 6). Infection typically initiates in mucosal epithelial cells and macrophages in lungs before disseminating to secondary lymphoid tissues, such as the spleen and peritoneal cavity (7, 8). Once in these compartments, the virus colonizes naïve B cells and drives a robust germinal center response (9, 10). This expansion of B cells is critical for the viral life cycle, allowing the virus to differentiate into memory B cells, the primary reservoir for lifelong latency, or plasma cells, from which the virus will reactivate from (11). Notably, the high proliferative rate of germinal center B cells, combined with the downregulation of tumor suppressors, makes this niche a primary site for cellular transformation, leading to the high incidence of germinal center-derived B-cell lymphomas associated with these infections (12-15). Despite their clinical significance, the host factors that facilitate the establishment of these chronic infections remain incompletely understood, largely due to the extreme species-specificity of human gammaherpesviruses which limits *in vivo* mechanistic studies. To overcome these limitations, murine gammaherpesvirus 68 (MHV68) is utilized as a tractable and biologically relevant animal model (5, 6). MHV68 shares significant genetic and pathogenetic homology with its human counterparts, including the ability to replicate in lung epithelium before establishing a robust latent reservoir in germinal center B cells (5, 16-18). In this model, the establishment of latency is a highly orchestrated process where the virus usurps host immune signaling to promote its own latent reservoir. Among the host factors potentially involved in this process, interleukin-22 (IL-22) emerges as a candidate of significant interest due to its potent regulatory effects on mucosal barriers and lymphoid architecture (19, 20).

Interleukin-22 (IL-22), originally named as IL-10-related T cell-derived inducible factor (IL-TIF), is a member of IL-10 family of cytokines (21, 22). IL-22 is expressed by various immune cell subsets, including activated natural killer (NK) cells, NKT cells, neutrophils, innate lymphoid cells (ILCs), γδ T cells, CD8+ T cells (23-25) and CD4+ T cell subsets including of the TH17 (26, 27), TH1 (28) and TH22 (29). Uniquely, IL-22 signaling is directed toward non-hematopoietic cells through a heterodimeric receptor complex (IL-22Rα and IL-10Rβ). IL-10Rβ is broadly expressed by most cell types, whereas the expression of IL-22Rα is restricted to non-hematopoietic cells (27, 30). IL-22 downstream signaling leads to the activation of Jak/STAT pathway—primarily STAT3—and MAP kinase/p38 pathways (30, 31). While IL-22 has a well-defined role in clearing bacterial and fungal pathogens, its role in viral infections is increasingly recognized as complex and context-dependent.

Emerging reports suggest that IL-22 can be either host-protective or proviral. In Rotavirus and MCMV infections, IL-22 exerts antiviral effects through cooperation with other cytokines or the recruitment of neutrophils (32, 33). Conversely, in West Nile Virus and Hepatitis B models, IL-22 can exacerbate tissue pathology or support virus-driven inflammation. This is achieved by inducing the expression of pro-inflammatory chemokines such as CXCL1, CXCL5, and CXCL10, which promotes the excessive recruitment of neutrophils and other inflammatory infiltrates into infected tissues (34, 35). Furthermore, EBV-induced infectious mononucleosis has been shown to elevate serum IL-22 levels in both adults and children (36, 37). Despite these insights, the specific role of host IL-22 in orchestrating the gammaherpesvirus life cycle and the establishment of latency remains unknown.

Our previous work identified a proviral role for the IL-17RA signaling axis during the establishment of chronic MHV68 infection (38-40). Given that IL-22 and IL-17 are often co-expressed and share functional synergy in inducing antimicrobial peptides (41), we hypothesized that IL-22 might have an influence on the MHV68 infection. Notably, gammaherpesviruses do not encode a discernable IL-22 homolog, they must, therefore, rely on the modulation of host-derived IL-22 to potentially influence the immunological niche required for latency.

In this study, we investigated the role of host IL-22 during MHV68 chronic infection. We report that MHV68 infection triggers a systemic and tissue-specific upregulation of IL-22, driven primarily by virus-specific CD4⁺ and CD8⁺ T cells. Contrary to its role in bacterial defense, we demonstrate that IL-22 serves as a proviral factor. Our data show that IL-22^-/-^ mice exhibit a reduction in the latent viral reservoir and a blunted germinal center response. Furthermore, we reveal that IL-22 is critical for the induction of the virus-mediated polyclonal B-cell response and the subsequent production of self-directed autoantibodies. These findings provide a novel paradigm for IL-22 biology, identifying it as a host resource co-opted by gammaherpesviruses to facilitate the chronic infection and virus-induced autoimmunity.

## Results

### MHV68 infection elicits IL-22 expression across multiple anatomic sites

Due to the extreme species-specificity of human gammaherpesviruses EBV and KSHV, *in vivo* studies of their host interactions are inherently limited. To address this, we utilized the genetically and biologically related murine gammaherpesvirus 68 (MHV68), an established and tractable model for studying gammaherpesvirus pathogenesis in an immunocompetent host (5, 42-44).

Because gammaherpesviruses do not encode a discernable IL-22 homolog, we first characterized the host IL-22 expression patterns in the lungs and spleen, the two primary sites of acute and chronic viral reservoirs. We examined C57BL/6 wild-type (BL6) mice at 9 days post-infection (dpi) and 16 dpi, representing the peaks of lytic and latent infection, respectively. Intranasal infection with MHV68 triggered a spike in IL-22 protein levels in the lungs at both 9 and 16 dpi (Fig. 1A, C). In the spleen, IL-22 levels were trending higher at 9 dpi, though they did not reach statistical significance (Fig. 1B). However, the onset of latent infection significantly upregulated splenic IL-22 levels by 16 dpi (Fig. 1D). Given that IL-22 is produced by a diverse array of immune cells, we investigated whether virus-specific T cells contribute to this response. Using the gating strategies defined in Fig. S1, we analyzed viral specific, splenic CD4+ and CD8+ T cell populations at 16 dpi. Following *ex vivo* restimulation with the immunodominant MHV68 peptides gp150 (MHC class II) and ORF6 (MHC class I), we observed a significant increase in the total numbers of IL-22-expressing, virus-specific CD4+ and CD8+ T cells (Fig. 1E-F). Furthermore, systemic IL-22 levels in the serum were significantly elevated at 16 dpi (Fig. 1G), indicating that MHV68 infection induces a systemic IL-22 response during the chronic phase.

**Fig 1.**
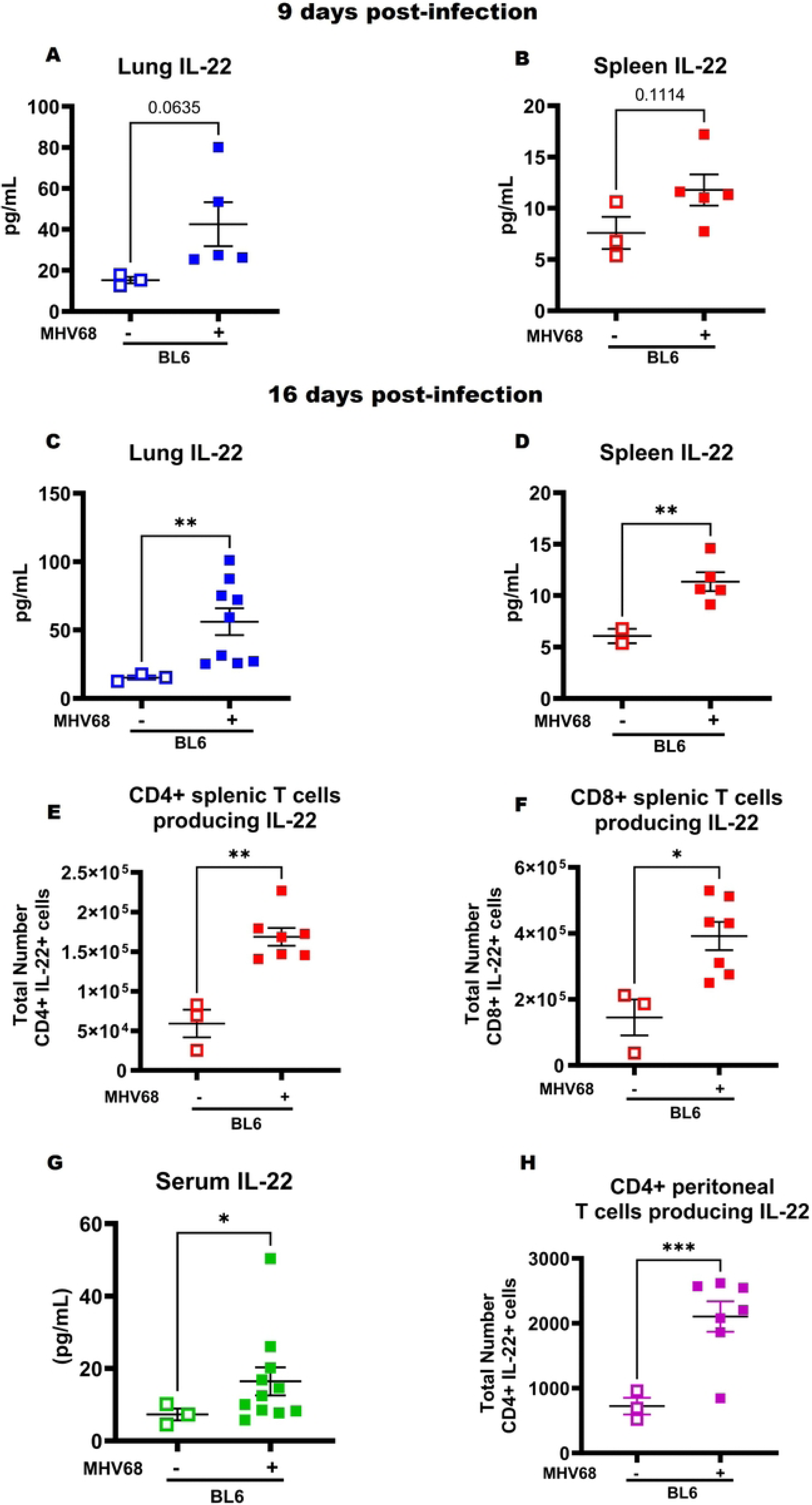
MHV68 infection triggers systemic and tissue-specific IL-22 production. C57BL/6 (BL6) mice were intranasally infected with 1000 PFU of MHV68. Tissues and serum were harvested at 9 dpi and 16 dpi to evaluate IL-22 expression during the transition from lytic to latent infection. (A–D) The respective IL-22 levels 9- and 16-days post-infection in the lungs (A, C) and spleens (B, D) were analyzed via ELISA. (E) Systemic IL-22 levels in the serum at 16 dpi was determined by a flow cytometry-based multiplex bead assay. (F, G) Total numbers of IL-22-expressing splenic CD4+ (F) and CD8+ (G) T cells at 16 dpi. Splenocytes were restimulated *ex vivo* with the MHV68 immunodominant peptides gp150 or ORF6, followed by intracellular cytokine staining (ICS) and flow cytometric analysis. (H) Total numbers of IL-22-expressing CD4+ T cells in the peritoneal cavity at 16 dpi as determined by flow cytometry. Each experimental group consists of 3–4 animals and each symbol represent an individual mouse. Data are pooled from two to three independent experiments. Mean and standard error of the mean are shown. \**P* < 0.05, \*\**P*< 0.01, ***P <.001

Finally, as the peritoneal cavity serves as another critical reservoir for latent MHV68 during chronic infection, we examined IL-22 expression in this compartment. We observed a significant increase in the total number of IL-22-expressing CD4+ T cells within the peritoneal cavity of MHV68-infected mice (Fig. 1H). Collectively, these data demonstrate that MHV68 infection elicits a robust IL-22 response across multiple anatomic locations during both the acute and chronic stages of infection.

### IL-22 expression supports gammaherpesvirus latency and ex-vivo reactivation

After observing the significant induction of IL-22 across multiple anatomic sites following MHV68 infection, we next investigated its functional role during chronic infection. (BL6) and IL-22-deficient (IL-22^-/-^) mice were infected intranasally with MHV68. We evaluated latency parameters at 16 dpi, the peak of chronic infection, within the spleen and peritoneal cavity, the two primary reservoirs for latent MHV68 (5).

To quantify the latent viral load, we performed limiting-dilution PCR (LD-PCR) analysis to determine the frequency of MHV68 DNA-positive cells. Compared to infected BL6 controls, IL-22^-/-^ mice exhibited a reduction in the frequency of MHV68 DNA-positive cells in both the spleen and the peritoneal cavity (Fig. 2A, C). Since gammaherpesviruses can periodically reactivate from the latent phase (45), we next assessed the impact of IL-22 deficiency on viral reactivation competency via limiting dilution assay (LDA). Explanted splenocytes and peritoneal cells were serially diluted onto mouse embryonic fibroblast (MEF) monolayers and monitored for cytopathic effects (CPE) at 21 days post-challenge. Consistent with the reduction in DNA-positive cells, the frequency of *ex-vivo* MHV68 reactivation was significantly lower in IL-22^-/-^ mice than in their BL6 counterparts at 16 dpi (Fig. 2B, D). These data demonstrate that IL-22 signaling supports both the establishment of peak latency and the capacity for viral reactivation.

**Figure 2.**
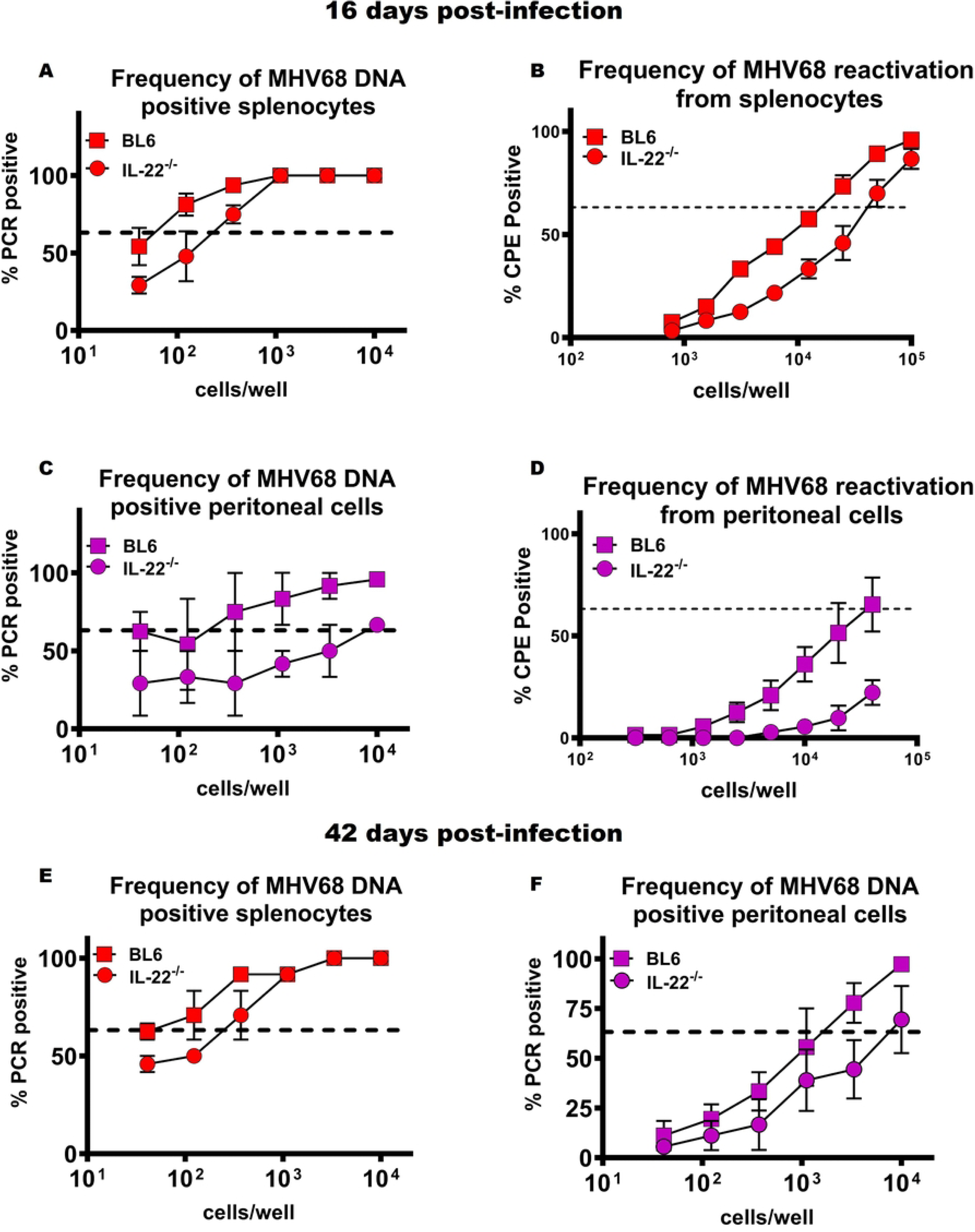
IL-22 deficiency reduces latent viral reservoir and *ex vivo* reactivation. BL6 and IL-22^-/-^ mice were infected as described in the legend to Fig 1. Splenocytes and peritoneal cells were harvested at 16 dpi (peak latency) and 42 dpi (long-term latency). (A, C) The frequency of MHV68 DNA-positive cells in the spleen (A) and peritoneal cavity (C) at 16 dpi were determined by limiting-dilution PCR (LD-PCR). (B, D) The frequency of *ex vivo* viral reactivation from splenocytes (B) and peritoneal cells (D) at 16 dpi. Cells were serially diluted onto MEF monolayers and monitored for cytopathic effect (CPE) at 21 days post-challenge. (E, F) The frequency of MHV68 DNA-positive cells in the spleen (E) and peritoneal cavity (F) during long-term infection (42 dpi) were determined by LD-PCR. Each experimental group consists of three to five animals. Data were pooled from two to five independent experiments, and SEM is displayed. In the limiting dilution assays, the dotted line is drawn at 63.2%, and the x-coordinate of the intersection of this line with the sigmoid graph represents the inverse of the frequency of positive events.

Following the 16-dpi peak, MHV68 latency typically begins to stabilize. By 42 dpi, the latent reservoir reaches a minimal "set-point" that is maintained for the life of the host, usually accompanied by a reduction in viral reactivation to levels below the limit of detection (5). We next evaluated whether IL-22 regulates this long-term phase of infection. Similar to the trends observed during peak latency at 16dpi, IL-22 deficiency resulted in a sustained reduction in the frequency of MHV68 DNA-positive cells in the spleen and peritoneal cavity at 42 dpi (Fig. 2E, F). As expected, no detectable reactivation was observed in either IL-22^-/-^ or BL6 mice at this late time point. These findings indicate that while IL-22 is essential for reaching peak levels of MHV68 latency, it also plays a sustained role in the maintenance of the long-term viral reservoir.

### Loss of IL-22 selectively attenuates splenic virus-specific CD8⁺ T Cell expansion

Having observed a significant reduction in viral latency and *ex vivo* reactivation in MHV68-infected IL22^-/-^ mice at the peak of latency (Fig. 2), we next investigated whether this diminished viral burden influenced the host immune response. CD8+ T cells are essential for controlling the latent reservoir and limiting viral reactivation through a dual strategy: first, viral specific CD8+ T cells prune back the initial surge of infected B cells (46). Second, they use perforin-mediated lysis and IFNγ secretion to ‘lock’ the viral genome in dormant state (47). Therefore, we quantified the frequency and absolute numbers of activated CD8+ T cells in the spleen and peritoneal cavity of infected BL6 and IL-22^-/-^ mice during chronic infection. While the global activation of the CD8+ T cells was comparable between groups (Fig. 3A-D), the frequency and absolute numbers of virus-specific CD8+ T cells were significantly reduced in the spleens of IL-22^-/-^ mice (Fig. 3 E-G). This reduction matches our finding of a reduced splenic viral load (Fig. 1A), suggesting that in absence of IL-22, there is not enough virus present to trigger the robust T-cell expansion typically seen in BL6 mice. Notably, this defect was restricted to the lymphoid compartment as the magnitude of the virus-specific CD8+ T cell response in the peritoneal cavity remained unchanged (Fig. 3H-J).

**Figure 3.**
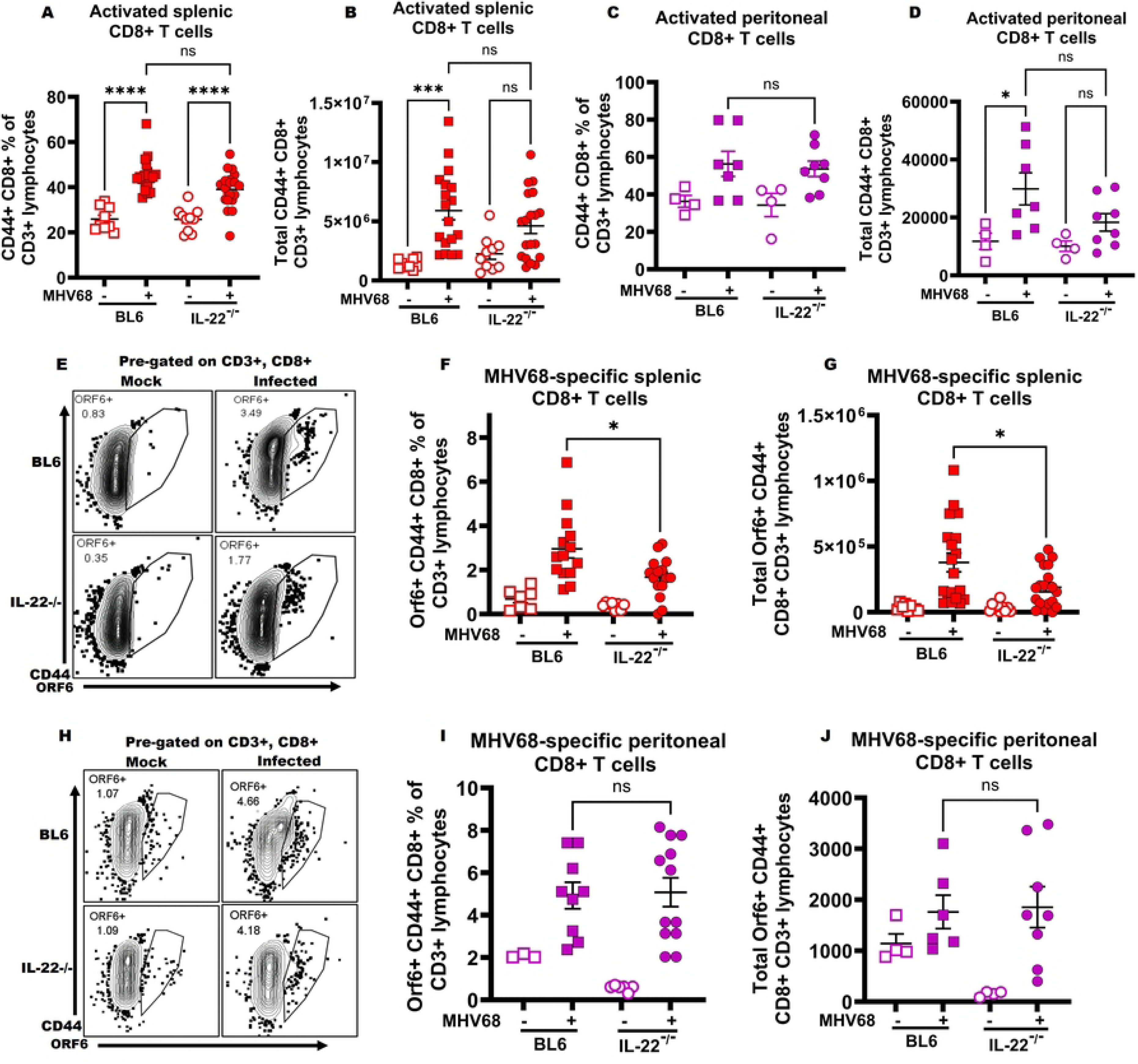
IL-22 deficiency selectively impairs splenic virus-specific CD8+ T cell expansion. WT (BL6) and IL-22^-/-^ mice were infected as described in the legend to Fig 1. CD8+ T cells were assessed at 16 dpi (peak latency) in the spleen and peritoneal cavity (PEC). (A–D) Frequency and absolute numbers of activated CD8+T cells (defined as CD3+CD8+ CD44+ cells) in the spleen (A, B) and PEC (C, D) were determined by flow cytometry. (E-J) Frequency and absolute number of MHV68-specific CD8+T cells (defined CD3+CD8+CD44+ORF6+) in the spleen (E-G) and peritoneal cavity (H-J), identified via MHC-I tetramer staining for the immunodominant ORF6_487-495_ (p56) epitope were determined by flowcytometry. Each experimental group consists of 3–4 animals and each symbol represent an individual mouse. Data are pooled from two to five independent experiments. Mean and standard error of the mean are shown. \**P* < 0.05, ***P <.001

### IL-22 is not required for CD8+ T cell functional capacity or systemic inflammatory stability

Because the reduction in virus-specific CD8+ T cells was restricted to spleen, we examined the functional capacity of these cells to produce gamma interferon (IFN-γ) in the absence of IL-22. Splenocytes were restimulated ex vivo with MHV68-derived peptide (ORF6_487-495_). The restimulation revealed that both the frequency and absolute numbers of IFNγ-producing splenic CD8+ T cells were equivalent between IL-22^-/-^ and BL6 mice (Fig. 4AI-C).

**Fig 4.**
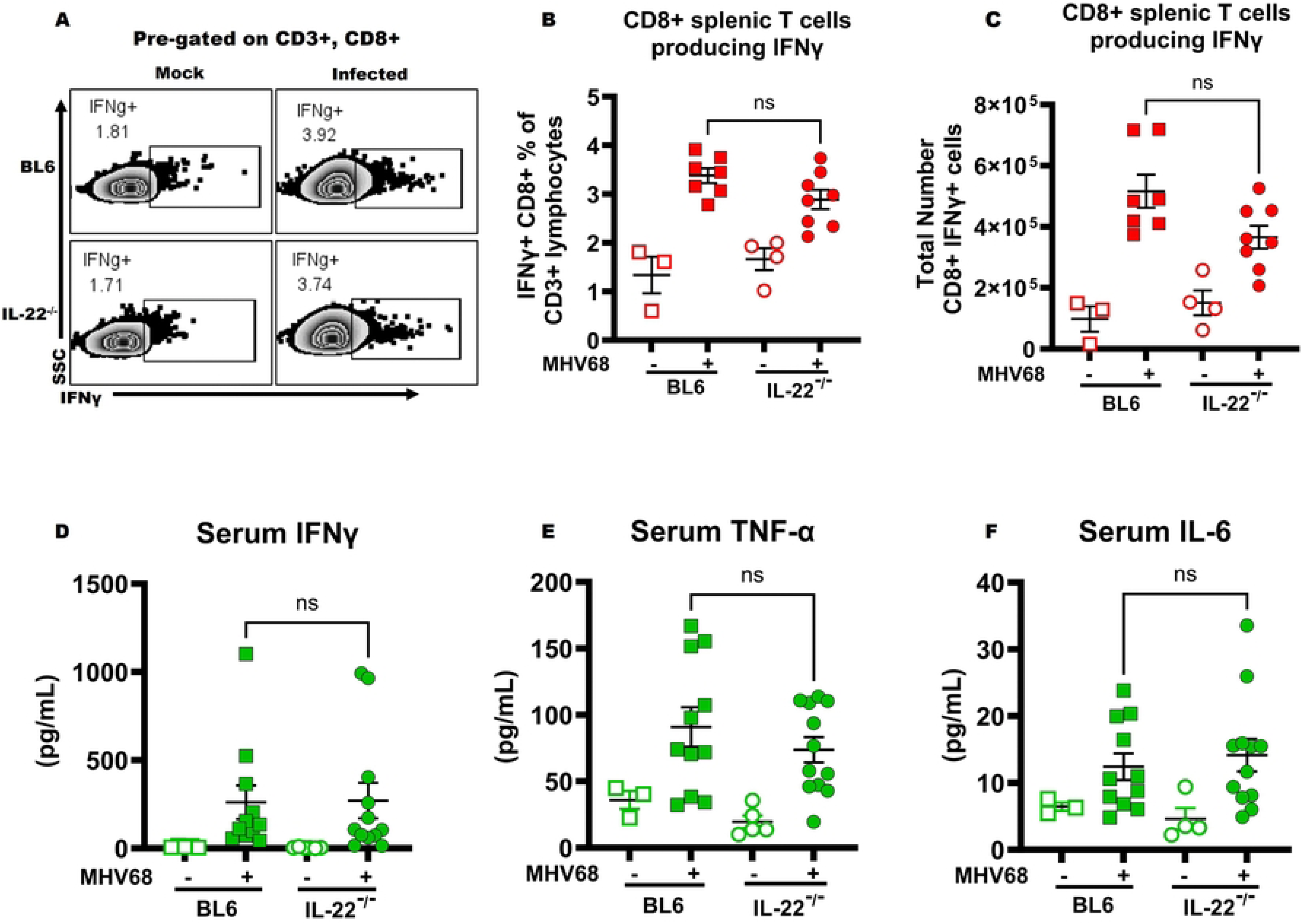
IL-22 is not required for CD8+ T cell functional capacity or systemic inflammatory stability. (A-C) Flow cytometry-based analysis of IFNγ production by viral specific splenic CD8+T cells. Splenocytes were restimulated *ex vivo* with Orf6 MHV68 peptide, and the frequency (A,B) and absolute number (C) of IFNγ+ cells were determined by intracellular cytokine staining (ICS). (D–F) Systemic inflammatory milieu at 16 dpi. Serum levels of IFNγ (D), TNFα (E), and IL-6 (F) were measured using a flow cytometry-based multiplex bead assay. Each experimental group consists of 3–4 animals and each symbol represent an individual mouse. Data are pooled from two to three independent experiments. Mean and standard error of the mean are shown.

Furthermore, the systemic inflammatory milieu remained stable, with no detectable changes in serum IFNγ, TNFα, or IL-6 levels (Fig. 4D-F). Together, these data suggest that the reduced splenic virus-specific CD8⁺ T cell response in IL-22^⁻/⁻^ mice reflects a lower antigenic burden rather than an intrinsic defect in CD8⁺ T cell effector function or a state of systemic inflammation in the absence of IL-22. These findings support a proviral role for IL-22 in the establishment of chronic gammaherpesvirus infection.

### IL-22 deficiency impairs germinal center expansion independent of global B cell activation and splenic pathology

A characteristic feature of MHV68 infection is the induction of splenomegaly, driven by the lymphoproliferative expansion of both B and T cells (48). Therefore, we next sought to determine if the loss of IL-22 altered the MHV68-induced splenic pathogenesis. Consistent with our findings that demonstrate intact global CD8+ T cell effector function and stable systemic cytokine levels (Fig. 4), IL-22^-/-^ mice developed gross splenomegaly indistinguishable from infected BL6 controls as the splenocytes expanded to a similar degree in both genotypes (Fig. 5A). The results indicate that IL-22 does not alter the MHV68-induced splenomegaly. We further dissected the splenic B cell compartment to determine if the reduction in viral latency in IL-22^-/-^ mice (Fig. 2) was a consequence of a global B cell defect. While MHV68 infection induced a robust increase in the absolute numbers of total B220+ B cells, no significant differences were observed between infected BL6 and IL-22^-/-^ mice (Fig. 5B). This suggests that the initial activation and splenic accumulation of B cells, the primary host for MHV68 latency, occur independently of IL-22 signaling. Notably, gammaherpesviruses specifically usurp germinal center B cell differentiation to establish long-term latent infection in memory B cells (49, 50). At the peak of MHV68 latency at 16 days post infection, the majority of latent viral reservoir is hosted by the germinal center B cells (49, 50), with CD4+ T follicular helper cells (Tfh) playing a critical role in the MHV68-driven germinal center response (51, 52). We next investigated whether IL-22 was required for the inducing MHV68-driven germinal center response. Strikingly, in contrast to the comparable total B cell numbers in the spleen (Fig. 5B), we observed a significant reduction in both the frequency and absolute number of germinal center B cells in IL-22^-/-^ mice at the peak of latency (Fig. 5C-E). Consistent with decreased germinal center B cells, MHV68-infected IL22^-/-^ mice also showed decreased proportion and absolute numbers of T follicular helper cells (Fig. 5F-H). Thus, IL-22 expression supported the MHV68-driven germinal center response during chronic infection.

**Figure 5.**
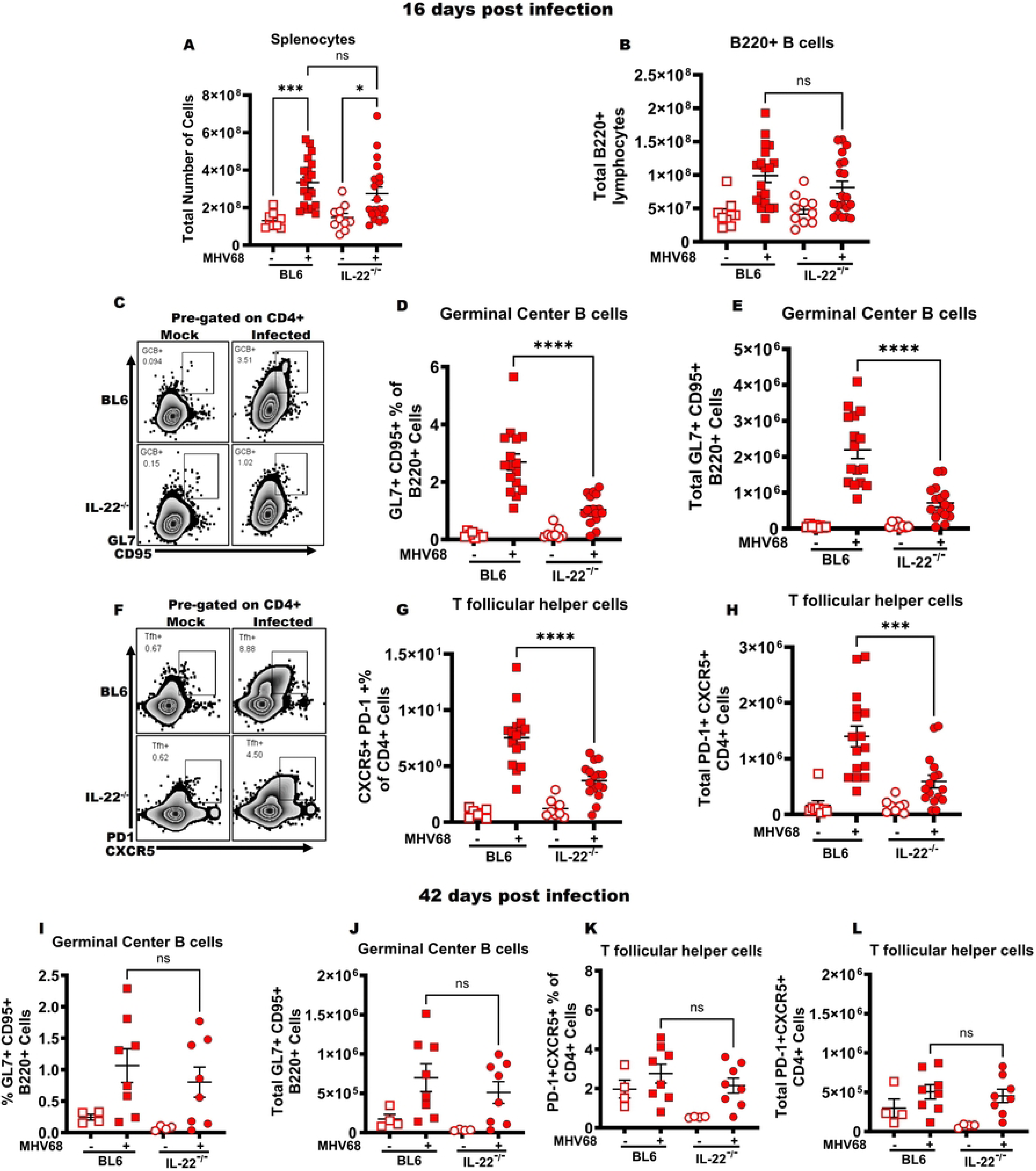
IL-22 deficiency impairs germinal center expansion independent of global B cell activation and splenic pathology. WT (BL6) and IL-22^-/-^ mice were infected as in Fig 1. Splenocytes were harvested at 16 dpi (peak latency) and 42 dpi (long-term latency) to evaluate splenomegaly and B cell differentiation. (A-B) Total numbers of splenocyte (A) and splenic-B cells (B220+) (B) at 16 dpi were determined by flow cytometry. The germinal center response was measured at 16- and 42-days post-infection in the spleen, with germinal center B cells (C-E; I,J) defined as B220+GL7+CD95+ cells and T follicular helper cells (F-H; K, L) defined as CD3+CD4+CXCR5+PD-1+cells. Each experimental group consists of 3–4 animals, and data are pooled from two to five independent experiments. Mean and standard error of the mean are shown. \**P* < 0.05, \*\**P* < 0.01, \*\*\*\**P* < 0.0001

To determine if IL-22^-/-^ deficiency sustains the reduction in the germinal center B cell at longer time points, we examined the splenic B cell compartment during long-term chronic infection (42 dpi). By this late stage of infection, the massive splenomegaly observed at 16dpi had largely resolved in both genotypes, with no significant differences in total numbers of splenocytes and B220+ B cells (Suppl. FigS2. A, B). Intriguingly, while the germinal center response was significantly blunted at the peak of latency, the frequency and absolute numbers of germinal center B cells were comparable between IL-22^-/-^ and BL6 mice at 42dpi (Fig. 5I-L). These results indicate that IL-22 supports the establishment of the virus-driven germinal center response but does not maintain long-term maintenance of germinal center B cells.

### IL-22 supports the production of irrelevant and self-directed class-switched antibodies stimulated by MHV68 infection

Having observed a significant reduction in the germinal center response in MHV68-infected IL-22^-/-^ mice (Fig. 5C-E), we next characterized the functional output of this impaired B cell compartment by measuring serum antibody titers. Gammaherpesviruses are unique in their ability to induce a robust, bystander (non-antigen-specific) B cell differentiation program. This leads to a rapid, albeit transient, increase in antibody titers against irrelevant and self-antigens (53), along with a delayed rise in virus-specific antibody titers (54). At 16 days post-infection, IL-22^-/-^ mice exhibited a significant reduction in total IgG and IgM titers compared to BL6 controls (Fig. 6A-B). Intriguingly, this defect was restricted to the polyclonal response, as MHV68-specific IgG and IgM titers remained comparable between IL-22^-/-^ and BL6 mice at the same time point (Fig. 6C-D).

**Figure 6.**
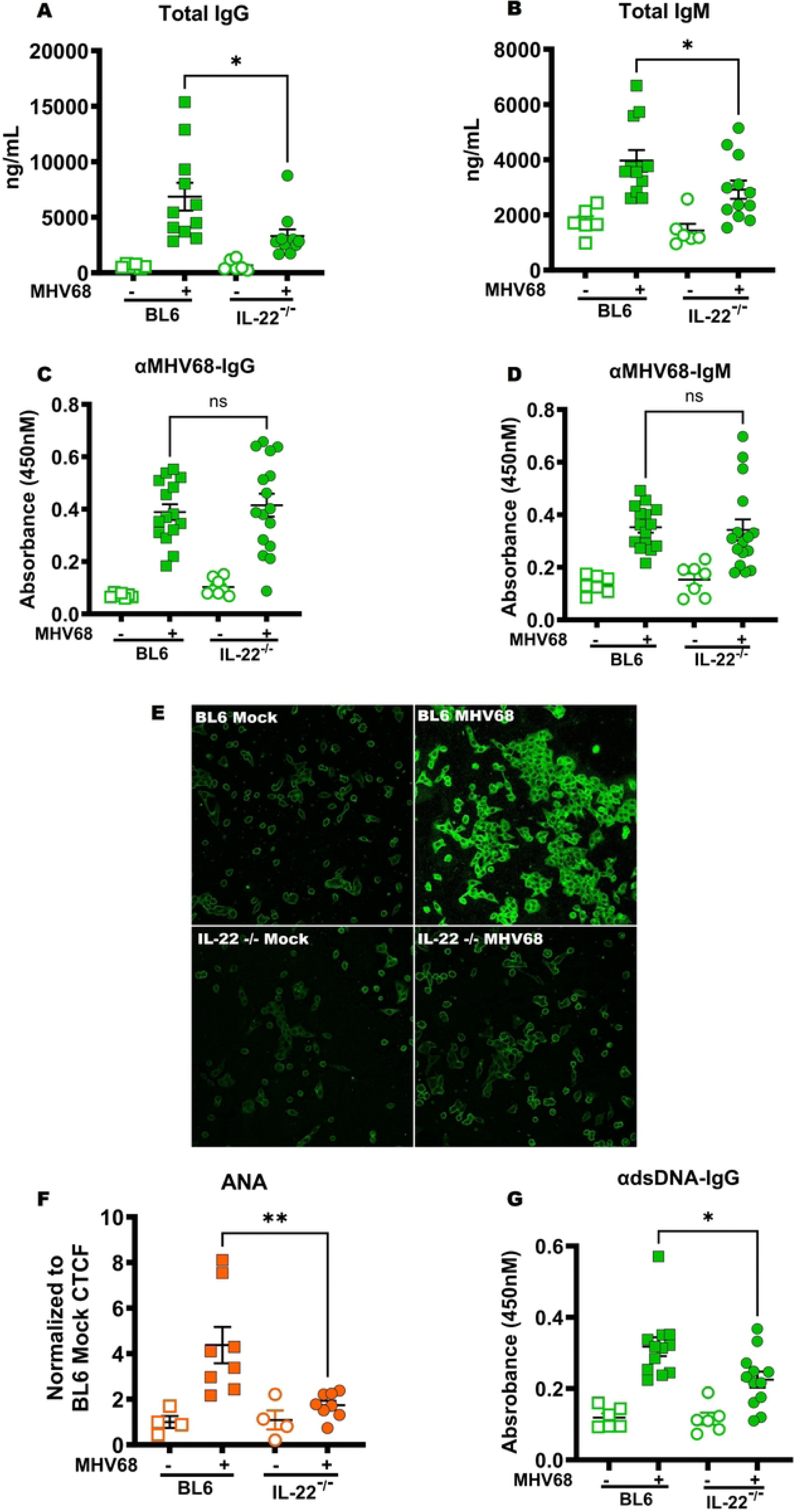
IL-22 deficiency reduced MHV68-induced polyclonal and autoantibody response. WT (BL6) and IL-22^-/-^ mice were either mock treated or infected as described in the legend to Fig 1. Sera were used to determine total IgG (A) and IgM (B), MHV68-specific IgG (C) and IgM (D), and dsDNA IgG (G) antibody titers via ELISA. (E) Reactivity of mouse sera with Hep-2 monolayers (ANA) using anti-mouse IgG fluorescein isothiocyanate (FITC)-conjugated antibody for detection (representative images), corrected total cell fluorescence (CTCF) quantified in panel (F). Data are pooled from two to five independent experiments, with each symbol representing an individual mouse. Mean and standard error of the mean are shown. \**P* < 0.05, \*\**P* < 0.01.

Gammaherpesviruses, including EBV and MHV68, induce a uniquely robust polyclonal increase in immunoglobulins directed against self-antigens and antigens of foreign species (53, 55). This induction of virus-nonspecific polyclonal antibody response forms the basis for the diagnostic assay, where high levels of antibodies directed against horse red blood cells are indicative of a recent EBV infection in humans (56). To capture the breadth of the self- and foreign-antigen-directed response in our model, we subjected sera from infected mice to an anti-nuclear antibody (ANA) assay, a standard clinical diagnostic for autoimmune reactivity (55). Consistent with our observations of reduced total immunoglobulins, MHV68-infected IL-22^-/-^ mice showed significantly diminished overall self-antigen reactivity compared to BL6 mice (Fig. 6E).

Furthermore, the qualitative nature of this autoantibody response was altered; the robust, pan-cellular staining pattern produced by sera from infected BL6 mice was markedly attenuated when using sera from IL-22^-/-^ mice, which displayed a significant decrease in nuclear staining intensity (Fig. 6F). To further define this loss of nuclear reactivity, we measured antibodies specific for double-stranded DNA (anti-dsDNA). While infected BL6 mice mounted a significant anti-dsDNA response, titers in IL-22^-/-^ mice were strikingly lower and exhibited a downward trend compared to infected BL6 mice (Fig. 6G). Collectively, these results demonstrate that IL-22 supports MHV68-induced expansion of the self-directed B cell repertoire and the production of "bystander" antibodies.

## Discussion

The hallmark of gammaherpesvirus infection is the establishment of a permanent latent reservoir (8). Using the MHV68 model, we demonstrated that the host cytokine IL-22 is a critical player in this process. Our results show that MHV68 infection triggers a robust increase in IL-22 across multiple tissues, driven primarily by virus-specific T cells. Crucially, we identified IL-22 as a proviral factor. In its absence, the latent viral reservoir is reduced, and the germinal center response is blunted. Furthermore, IL-22 supports polyclonal autoimmunity typically seen in these infections. Collectively, these findings reveal that the virus co-opts the host’s IL-22 response to build a protective niche for its own survival.

Our findings expand the understanding of IL-22 by identifying its previously unknown proviral role during gammaherpesvirus infection. While IL-22 is traditionally recognized for its host-protective and regenerative functions at mucosal barriers, particularly against bacterial and fungal pathogens, its role is increasingly understood to be highly context-dependent (57).

A major contribution of this study is the demonstration that gammaherpesviruses have evolved to exploit the IL-22 axis to facilitate chronic infection. While existing literature has explored the dual nature of IL-22 in various inflammatory contexts, no prior studies have investigated its role in gammaherpesvirus infection. Our data provide the first evidence that, in the context of these DNA viruses, IL-22 shifts from a barrier-protective factor to a facilitator of viral latency and reactivation.

### Mapping the IL-22 Response Over Time

Our study demonstrates that MHV68 infection triggers a significant increase in IL-22 levels within the lungs, spleen, and serum (Fig. 1). While IL-22 induction is a hallmark of the host response to diverse pathogens, ranging from bacterial infections like *Citrobacter rodentium* (58) *and Klebsiella pneumoniae (59)* to respiratory viruses like Influenza(60, 61), the temporal and spatial distribution we observed in this model is particularly striking. We identified a clear temporal shift in IL-22 expression: lung and splenic levels were trending high during the early acute phase, which reached to the significant levels during the chronic stage at 16 days post-infection. Furthermore, systemic IL-22 levels were also significantly higher at 16 days post-infection (Fig. 1G)

In many acute mucosal infections, innate lymphoid cells (ILCs), particularly ILC3s, serve as the rapid, primary source of IL-22 to promote epithelial barrier integrity (62, 63). While our study focused on the adaptive arm and identified virus-specific CD4+ and CD8+ T cells as major producers of IL-22 during the chronic phase of MHV68 infection, the potential contribution of ILCs, especially during the early acute peak in the lungs, remains an important area for future investigation. Regardless of the initial cellular source, the late-stage T-cell-derived IL-22 creates a paradoxical scenario: the adaptive immune response, typically intended to control the pathogen, instead provides the specific cytokine environment the virus needs to thrive. We propose that this represents a sophisticated form of evolutionary subversion, where MHV68 exploits a host pathway normally reserved for tissue repair and lymphoid organization. Although gammaherpesviruses do not encode their own IL-22 homolog, the virus effectively orchestrates a host environment rich in this cytokine, turning a host-protective "shield" into a proviral niche that facilitates a stable, lifelong latent reservoir. Further work is required to fully elucidate the molecular signaling pathways by which IL-22 influences B cell fate during infection, and to determine if this mechanism is conserved across other members including human pathogens like EBV and KSHV. Understanding these mechanisms will be critical in determining whether the IL-22 axis can be targeted to reduce the latent viral burden without compromising global antiviral immunity.

### How IL-22 Sustains Chronic Infection

Our data shows that IL-22 is required to both start and maintain the latent viral reservoir and reactivation (Fig. 2). Mice lacking IL-22 showed a significant reduction in viral DNA in the spleen and peritoneal cavity. These mice also showed less viral reactivation, meaning the virus struggled to exit a quiescent state and replicate without IL-22 signaling. This impact is long-lasting; even 42 days after infection, IL-22^-/-^ mice maintained significantly lower latent viral levels compared to wild-type controls. These findings suggest that IL-22 is a critical host factor that supports the long-term stability of the latent reservoir. By promoting a favorable microenvironment, IL-22 signaling ensures the virus can achieve and maintain its characteristic latent set-point throughout the chronic phase of infection. Notably, the mechanism by which IL-22 influences the germinal center response likely involves an indirect signaling axis. It is well-established that hematopoietic cells, including B and T cells, lack the functional IL-22 receptor (IL-22 Rα) (64). Therefore, the blunted germinal center response in IL-22^-/-^ mice is not a result of lost direct signaling to B cells. This suggests that IL-22 may instead act on non-immune populations within the lymphoid microenvironment to facilitate the expansion of the germinal center. These cells provide a cellular sanctuary that allows the virus to evade immune detection. This creates a proviral feedback loop: the virus triggers T cells to produce specific cytokines (like IL-22) which, instead of clearing the infection, actually help the virus-infected cells survive and expand. By exploiting this immune response, the virus secures its long-term establishment within the host reservoir.

### IL-22 Signaling Promotes a Polyclonal, Non-Specific Humoral Response

Beyond supporting the viral reservoir, IL-22 contributes to the unique immunopathology associated with gammaherpesvirus infection. Both EBV and MHV68 drive a massive, non-specific expansion of B cells by hijacking the germinal center reaction (53, 65). This process triggers a "polyclonal activation", where B cells are turned on regardless of their target, leading to a massive surge in total antibodies and the secretion of high levels of irrelevant and self-directed autoantibodies (53, 65). In IL-22^-/-^ mice, this non-specific humoral response was significantly dampened (Fig. 6E,F). Notably, this was not due to a global B cell defect, as the production of virus-specific antibodies remained intact (Fig. 6C,D). Instead, IL-22 signaling appears to lower the threshold for B cell selection that allows irrelevant B cells to bypass the usual quality control checks of the germinal center for high affinity antibody production (66). Our data suggest that while IL-22 is not necessary for a focused, virus-specific response (Fig. 6C,D), it is a key driver for the expansion of irrelevant B cell clones (Fig. 6E-f). This provides a mechanistic link between IL-22-mediated viral expansion and the transient loss of B cell tolerance typically observed during chronic gammaherpesvirus infection.

Gammaherpesviruses are unique in their ability to induce a robust germinal center response, which supports a majority of the latent viral reservoir (49, 53, 67). These viruses also drive an increase in polyclonal antibodies directed against non-relevant foreign antigens and even self-directed antigens (67, 68). Our study identifies the IL-22/ germinal center axis as a primary driver of gammaherpesvirus pathogenesis. By demonstrating that IL-22 supports peak viral latency (Fig. 2), the virus-driven germinal center response (Fig. 5), and the production of a diverse, non-specific antibody repertoire (Fig. 6), we provide a new framework for understanding how these viruses subvert host defenses to establish a long-term reservoir. While we have established the requirement for this pathway *in vivo*, the specific cellular targets of the IL-22 signal remain an intriguing question. Given that IL-22Rα is absent on hematopoietic cells, our findings strongly suggest that IL-22 does not communicate directly with the B or T cells it ultimately influences. Instead, the IL-22 signal must be integrated by non-immune populations within the germinal center microenvironment that supports differentiation of memory B cells, the long-term latent reservoir of the virus. Future research utilizing cell-specific deletions of the IL-22Rα complex will be essential to precisely map this indirect signaling circuit. Previous studies have indicated that IL-22 signaling can bolster the lymphoid architecture and support germinal center integrity in other contexts (20). Identifying the specific stromal or epithelial cells within the lymphoid architecture will clarify how IL-22-dependent changes in the microenvironment translate into the expansion of the germinal center response and the lowered threshold for B cell selection observed during infection.

Finally, the striking reduction in autoantibodies in IL-22^-/-^ mice suggests that the IL-22 axis may be a viable therapeutic target specifically in the context of gammaherpesvirus-driven immunopathology. If blocking IL-22 can disrupt the supportive niche for self-reactive B cells without compromising global antiviral immunity, it could represent a novel strategy for mitigating EBV-related autoimmune flares, such as those seen in Systemic Lupus Erythematosus (SLE) or Multiple Sclerosis (MS) (69). Understanding these nuances will be key to developing next-generation treatments that limit the establishment of chronic viral reservoir and mitigate its long-term damage to human health.

## Acknowledgements

This study was supported by startup funds provided to C.N.J. from Western Michigan University Homer Stryker M.D. School of Medicine. We thank the NIH Tetramer Core Facility (NIH Contract 75N93020D00005 and RRID:SCR_026557) for providing the MHV68 Orf6 tetramer used in this study.

## Author contributions

**S.T. Majeed:** Investigation, Formal analysis, Visualization, Validation, and Writing – Original Draft Preparation. **S.M. Bradford:** Investigation, Formal analysis, and Visualization. **C.N. Jondle:** Conceptualization, Investigation, Formal analysis, Visualization, Validation, Data curation, Funding acquisition, Supervision, Project administration, and Writing – Review & Editing.

**Declarations of interests.** The authors declare no competing interests.

## Materials and Methods

### Animal studies

All experimental manipulations of mice were approved by the Institutional Animal Care and Use Committee of Western Michigan University Homer Stryker M.D. School of Medicine (2022-0026). C57BL/6J and IL-22^-/-^ (C57BL/6-*Il22^tm1.1(icre)Stck^*/J, stock # 027524) mice were ordered from The Jackson Laboratories (Bar Harbor, ME). All mice were housed and bred in a specific-pathogen-free facility. Both male and female mice were used with no gender-specific phenotypes noted.

### Infections

MHV68 viral stock was prepared, and the virus titers on NIH 3T12 cells were determined. Six to ten-week-old mice were subjected to light anesthesia and intranasally inoculated with 1,000 PFU of MHV68 diluted in 15 µl sterile serum-free Dulbecco’s modified Eagle’s medium (DMEM). The spleen, lung, and peritoneal cells were harvested from euthanized mock-treated and MHV68-infected animals at indicated times postinfection. Mice were humanely euthanized using CO2 inhalation. Mice were bled before euthanasia via submandibular vein sampling, and the serum was collected using BD Microtainer blood collection tubes (Becton, Dickinson and Company, Franklin Lakes, NJ).

### Limiting dilution assays

The frequency of virally infected cells (cells harboring viral DNA) was determined by limiting-dilution (LD) PCR analysis, while the frequency of ex vivo reactivation to identify cells capable of producing infectious virus was determined by limiting-dilution-assay.

Spleens from MHV68 infected C57BL/6 and IL-22^-/-^ mice were mashed through 100-micron cell strainer (Fischer). Red blood cells were lysed by incubating in 1-2 ml of RBC lysis buffer (Sigma) for 3 minutes at 37°C. RBC lysis was neutralized by adding 18 mL DMEM and followed by 10-minute centrifugation at 1000rpm. The supernatant was discarded and splenocytes were resuspended in 12-16 ml of serum supplemented DMEM depending upon the pellet size.

Splenocytes were then filtered through 100-micron strainer (Fischer) to reduce clumping. Peritoneal lavage cells were isolated in 10ml of DMEM. The cells were centrifuged at 100rpm for 10 minutes. The supernatant was discarded and cells re-suspended in 0.5-3ml of serum supplemented DMEM depending upon the pellet size. Limiting-dilution (LD)-PCR analysis were performed as previously described (70). Briefly, splenocytes and peritoneal cells were pooled from all infected mice in each experimental group (three to five mice/group) and six threefold serial dilutions of latently infected cells were carried out in the background of uninfected 3T12 fibroblasts. Twelve replicates per dilution were loaded in 96 well PCR plate. The cells were digested overnight in presence of proteinase K and then subjected nested PCR using specific primers against the viral genome. Single-copy sensitivity and the absence of false-positive amplicons were confirmed using control standards. The samples were run on ethidium bromide stained 2% agarose gel for quantitation.

Ex-vivo reactivation efficiency was determined as previously described (70). Briefly, eight 2-fold serial dilutions of splenocytes or peritoneal cells from infected mice were plated on mouse embryonic fibroblasts (MEF) monolayers at 24 replicates per dilution. The presence of preformed infectious virus was detected by plating four 2-fold serial dilutions of mechanically disrupted splenocytes or peritoneal cells on MEF monolayers in parallel. Viral reactivation as indicated by the cytopathic clearing of MEFs was assessed on day 21 of culture. All replicates were scored in a binary fashion for the presence of live fibroblasts (no viral reactivation/replication) or their absence (cytopathic effect driven by lytic replication).

### Flow Cytometry

Single-cell suspensions of splenocytes and peritoneal cells from individual mice were prepared in fluorescence-activated cell sorting (FACS) buffer (phosphate-buffered saline, 2% fetal bovine serum, 0.05% sodium azide). 2x10^6^ cells were treated with Fc block (24G2) before extracellular staining for 30 min on ice. Data were acquired using an Attune NxT flowcytometer (Thermo Fisher Scientific, Waltham, MA) and analyzed using FlowJo software (Becton, Dickinson & Company, Ashland, OR). Following antibodies used in the study were purchased from BioLegend (San Diego, CA): CD3 (17A2), CD4 (RM4-5), CD8a (53-7.3), CD95 (Jo2), PD-1 (29f.1A12), B220 (RA3-6B2), GL7 (GL-7), CD44 (IM7), IL-22 (IL22JOP) and IFN-g (XMG1.2). CXCR5 (2G8) was purchased from BD Pharmingen (San Jose, CA). MHV68 Orf6 tetramer was obtained through the NIH Tetramer core (Atlanta, GA). Compensation controls were done using OneComp eBeads (Thermo Fisher Scientific, Waltham, MA). Briefly, negative control (beads alone) were used to establish a baseline PMT (photomultiplier tube) voltage and fluorescent background. Positive controls for each fluorochrome (beads with a single fluorochrome) were used to establish spill-over of the individual fluorochrome into the other channels being used. PMT values are adjusted for each fluorochrome to minimize spillover.

### T-cell phenotyping

For T-cell phenotyping, single-cell suspensions of splenocytes and peritoneal cells were plated at 4x10^6^ nucleated cells/mL in a 96-well plate. Non-specific stimulation of the cells was carried out using 10 ng/mL of phorbol 12-myristate 13-acetate (PMA; P8139-5MG; Sigma-Aldrich, St. Louis, MO), 1 mg/mL of ionomycin (I0634-1MG; Sigma-Aldrich), and 10 mg/mL of brefeldin A (420601; BioLegend, San Diego, CA) in DMEM with 10% FBS for 4 h at 37°C. For virus-specific CD4+ T-cell stimulation, 2.5 mg/mL of MHV68-specific viral peptide GP150 (71) (GenScript, Piscataway, NJ), and for virus-specific CD8+ T-cell stimulation 10mg/mL of ORF6 and ORF61 viral peptides (Thermo Fisher Scientific, Waltham, MA) were used in conjugation with 10mg/mL of brefeldin A (420601; BioLegend) in DMEM with 10% FBS for 6 h at 37°C. Following re-stimulation, cells were washed in FACS buffer before being treated with Fc block (24G2) and subjected to extracellular staining with an optimal antibody concentration for 30 min on ice. After extracellular staining, the cells were fixed and permeabilized using the BD Cytofix/Cytoperm kit (554714; Fisher Scientific, Hampton, NH). The cells were then intracellularly stained with an optimal antibody concentration for 30 min on ice. Data were acquired using Attune flow cytometer (Thermo Fisher) and analyzed using FlowJo software (Becton, Dickinson & Company, Ashland, OR).

### Ezymed-linked immunosobent assay (ELISA)

Total, MHV68-specific, and dsDNA immunoglobulin levels were determined as previously described (72). Briefly, Nunc Maxisorp plates (Fisher Scientific, Pittsburg, PA) were coated with either anti-IgG (heavy and light) or anti-IgM antibodies (Jackson ImmunoResearch, West Grove, PA), UV-irradiated MHV68 virus stock in PBS (740,000 micro joules/cm^2^X2) (Stratalinker UV Crosslinker 1800; Agilent Technologies, Santa Clara, CA), or dsDNA from Escherichia coli (12.5mg/ml; Sigma-Aldrich, St. Louis, MO) overnight at 4°C. Plates were washed with PBS-Tween (0.05%) and blocked for 1h with PBS-Tween (0.05%)-BSA (3%). This was followed by incubation with fivefold serial dilutions of serum in PBS-Tween (0.05%)-BSA (1.5%) for 2h and then washing with PBS-Tween (0.05%). Horseradish peroxidase (HRP)-conjugated goat anti-mouse total IgG (heavy and light chain [H1L]) or IgM (Jackson ImmunoResearch, West Grove, PA) along with 3,39,5,59-tetramethylbenzidine substrate (Life Technologies, Gaithersburg, MD) was used to detect the bound antibody. HRP enzymatic activity was stopped by the addition of 1 N HCl (Sigma-Aldrich, St. Louis, MO), and the absorbance was read at 450nm on BioTek EPOCH2 Plate Reader (Agilent Technologies, Santa Clara, CA).

### Muli-Analyte Flow Assay

Quantification of soluble IL-22, TNFα, IFNγ, and IL-6 concentrations in the serum were measured via LEGENDplex Mouse Th17 Panel (7-plex) multi-analyte flow assay kit (Cat# 741048) BioLegend (San Diego, CA) according to manufacturer directions and analyzed on AURORA spectral cytometer (Cytek, San Diego, CA). Data was processed with the LEGENDplex online data analysis software to determine the concentrations of the samples

### ANA panels

Antinuclear antibodies (ANAs) were assessed with an (ANA) test kit (Antibodies Inc., Davis, CA). Following the manufacturer’s protocol, serum was diluted (1:40 in PBS) and incubated over slides coated with fixed HEp-2 cells. Following serum incubation, the slides were rinsed and stained with anti-mouse IgG labeled with Alexa Fluor 488 (H+L) (ThermoScientific, Waltham, MA). Fluorescent images were captured using NIS Elements software. Corrected fluorescence was quantified using ImageJ software from a randomly chosen field of ∼ 20 cells in each sample.

### Protein extraction from tissues

For IL-22 detection from tissues, lung and spleen were harvested at the indicated time points post-infection. The tissues were disrupted in NP-40 lysis buffer (Thermo Scientific) and protease inhibitor cocktail (Thermo Scientific) by bead-beating using 1-mm zirconia/silica beads (Biospec Products, Bartelsville, OK). Homogenates were first incubated on ice for 20 minutes to allow complete lysis, followed by centrifugation at 20,000×g for 20 min. The supernatants were collected, and protein concentrations measured with a protein assay kit (Thermo Scientific). Equal amounts of proteins (100µg) were loaded for IL-22 detection using IL-22 enzyme-limited immunosorbent assay (ELISA) Max Deluxe set (BioLegend, San Diego, CA) according to the manufacturer’s instructions, and Nunc Maxi Sorp flat-bottomed plates (Thermo Fisher Scientific, Waltham, MA).

### Statistical analyses

Statistical analyses were performed using Student’s *t* test when comparing two groups and one-way ANOVA with Tukey’s post hoc test when comparing more than two groups (Prism, GraphPadSoftware, Inc.).

## Notes

### Competing Interest Statement

The authors have declared no competing interest.

## References

1. Wilson SJ, Tsao EH, Webb BL, Ye H, Dalton-Griffin L, Tsantoulas C, et al. X box binding protein XBP-1s transactivates the Kaposi’s sarcoma-associated herpesvirus (KSHV) ORF50 promoter, linking plasma cell differentiation to KSHV reactivation from latency. Journal of virology. 2007;81(24):13578–86.

2. Cesarman E. Gammaherpesviruses and lymphoproliferative disorders. Annual review of pathology. 2014;9:349–72.

3. Jha HC, Banerjee S, Robertson ES. The Role of Gammaherpesviruses in Cancer Pathogenesis. Pathogens (Basel, Switzerland). 2016;5(1).

4. Bjornevik K, Cortese M, Healy BC, Kuhle J, Mina MJ, Leng Y, et al. Longitudinal analysis reveals high prevalence of Epstein-Barr virus associated with multiple sclerosis. Science. 2022;375(6578):296–301.

5. Barton E, Mandal P, Speck SH. Pathogenesis and host control of gammaherpesviruses: lessons from the mouse. Annual review of immunology. 2011;29:351–97.

6. Wang Y, Tibbetts SA, Krug LT. Conquering the Host: Determinants of Pathogenesis Learned from Murine Gammaherpesvirus 68. Annual review of virology. 2021;8(1):349–71.

7. Gaspar M, May JS, Sukla S, Frederico B, Gill MB, Smith CM, et al. Murid herpesvirus-4 exploits dendritic cells to infect B cells. PLoS pathogens. 2011;7(11):e1002346.

8. Majeed ST, Jondle CN. Bringing Balance: Immune Interactions Regulating Murine Gammaherpesvirus 68 Latency. Current Clinical Microbiology Reports. 2024;11(1):1–11.

9. Flano E, Kim IJ, Woodland DL, Blackman MA. Gamma-herpesvirus latency is preferentially maintained in splenic germinal center and memory B cells. The Journal of experimental medicine. 2002;196(10):1363–72.

10. Roughan JE, Thorley-Lawson DA. The intersection of Epstein-Barr virus with the germinal center. Journal of virology. 2009;83(8):3968–76.

11. Stebegg M, Kumar SD, Silva-Cayetano A, Fonseca VR, Linterman MA, Graca L. Regulation of the Germinal Center Response. Frontiers in Immunology. 2018;9(2469).

12. Phan RT, Dalla-Favera R. The BCL6 proto-oncogene suppresses p53 expression in germinal-centre B cells. Nature. 2004;432(7017):635–9.

13. Muramatsu M, Sankaranand VS, Anant S, Sugai M, Kinoshita K, Davidson NO, et al. Specific expression of activation-induced cytidine deaminase (AID), a novel member of the RNA-editing deaminase family in germinal center B cells. The Journal of biological chemistry. 1999;274(26):18470–6.

14. Martin A, Bardwell PD, Woo CJ, Fan M, Shulman MJ, Scharff MD. Activation-induced cytidine deaminase turns on somatic hypermutation in hybridomas. Nature. 2002;415(6873):802–6.

15. Küppers R. B cells under influence: transformation of B cells by Epstein-Barr virus. Nat Rev Immunol. 2003;3(10):801–12.

16. Sunil-Chandra NP, Efstathiou S, Nash AA. Murine gammaherpesvirus 68 establishes a latent infection in mouse B lymphocytes in vivo. J Gen Virol. 1992;73 (Pt 12):3275–9.

17. Collins CM, Speck SH. Tracking murine gammaherpesvirus 68 infection of germinal center B cells in vivo. PLoS One. 2012;7(3):e33230.

18. Stewart JP, Usherwood EJ, Ross A, Dyson H, Nash T. Lung epithelial cells are a major site of murine gammaherpesvirus persistence. J Exp Med. 1998;187(12):1941–51.

19. Wei HX, Wang B, Li B. IL-10 and IL-22 in Mucosal Immunity: Driving Protection and Pathology. Front Immunol. 2020;11:1315.

20. Barone F, Nayar S, Campos J, Cloake T, Withers DR, Toellner KM, et al. IL-22 regulates lymphoid chemokine production and assembly of tertiary lymphoid organs. Proceedings of the National Academy of Sciences of the United States of America. 2015;112(35):11024–9.

21. Dumoutier L, Louahed J, Renauld JC. Cloning and characterization of IL-10-related T cell-derived inducible factor (IL-TIF), a novel cytokine structurally related to IL-10 and inducible by IL-9. J Immunol. 2000;164(4):1814–9.

22. Dumoutier L, Van Roost E, Colau D, Renauld JC. Human interleukin-10-related T cell-derived inducible factor: molecular cloning and functional characterization as an hepatocyte-stimulating factor. Proceedings of the National Academy of Sciences of the United States of America. 2000;97(18):10144–9.

23. Wolk K, Kunz S, Asadullah K, Sabat R. Cutting edge: immune cells as sources and targets of the IL-10 family members? Journal of immunology (Baltimore, Md : 1950). 2002;168(11):5397–402.

24. Goto M, Murakawa M, Kadoshima-Yamaoka K, Tanaka Y, Nagahira K, Fukuda Y, et al. Murine NKT cells produce Th17 cytokine interleukin-22. Cell Immunol. 2009;254(2):81–4.

25. Moreira-Teixeira L, Resende M, Coffre M, Devergne O, Herbeuval JP, Hermine O, et al. Proinflammatory environment dictates the IL-17-producing capacity of human invariant NKT cells. Journal of immunology (Baltimore, Md : 1950). 2011;186(10):5758–65.

26. Liang SC, Tan XY, Luxenberg DP, Karim R, Dunussi-Joannopoulos K, Collins M, et al. Interleukin (IL)-22 and IL-17 are coexpressed by Th17 cells and cooperatively enhance expression of antimicrobial peptides. J Exp Med. 2006;203(10):2271–9.

27. Zheng Y, Danilenko DM, Valdez P, Kasman I, Eastham-Anderson J, Wu J, et al. Interleukin-22, a T(H)17 cytokine, mediates IL-23-induced dermal inflammation and acanthosis. Nature. 2007;445(7128):648–51.

28. Volpe E, Touzot M, Servant N, Marloie-Provost MA, Hupé P, Barillot E, et al. Multiparametric analysis of cytokine-driven human Th17 differentiation reveals a differential regulation of IL-17 and IL-22 production. Blood. 2009;114(17):3610–4.

29. Eyerich S, Eyerich K, Pennino D, Carbone T, Nasorri F, Pallotta S, et al. Th22 cells represent a distinct human T cell subset involved in epidermal immunity and remodeling. J Clin Invest. 2009;119(12):3573–85.

30. Wolk K, Kunz S, Witte E, Friedrich M, Asadullah K, Sabat R. IL-22 increases the innate immunity of tissues. Immunity. 2004;21(2):241–54.

31. Liang SC, Nickerson-Nutter C, Pittman DD, Carrier Y, Goodwin DG, Shields KM, et al. IL-22 induces an acute-phase response. J Immunol. 2010;185(9):5531–8.

32. Stacey MA, Marsden M, Pham NT, Clare S, Dolton G, Stack G, et al. Neutrophils recruited by IL-22 in peripheral tissues function as TRAIL-dependent antiviral effectors against MCMV. Cell Host Microbe. 2014;15(4):471–83.

33. Hernández PP, Mahlakoiv T, Yang I, Schwierzeck V, Nguyen N, Guendel F, et al. Interferon-λ and interleukin 22 act synergistically for the induction of interferon-stimulated genes and control of rotavirus infection. Nat Immunol. 2015;16(7):698–707.

34. Wang P, Bai F, Zenewicz LA, Dai J, Gate D, Cheng G, et al. IL-22 signaling contributes to West Nile encephalitis pathogenesis. PLoS One. 2012;7(8):e44153.

35. Zhang Y, Cobleigh MA, Lian JQ, Huang CX, Booth CJ, Bai XF, et al. A proinflammatory role for interleukin-22 in the immune response to hepatitis B virus. Gastroenterology. 2011;141(5):1897–906.

36. Li D, Mao K, Luo P, Zheng Z, Liu J, Zhang C, et al. Elevated interleukin-10, -22, -24, and -26 in serum samples of children with infectious mononucleosis. J Med Biochem. 2025;44(4):776–83.

37. Li Y, Li LX, Gao Y. [The Role of Notch Signaling Pathway in Adult Patients with Epstein-Barr Virus-induced Infectious Mononucleosis]. Zhongguo Shi Yan Xue Ye Xue Za Zhi. 2024;32(3):920–6.

38. Jondle CN, Johnson KE, Aurubin C, Sylvester P, Xin G, Cui W, et al. Gammaherpesvirus Usurps Host IL-17 Signaling To Support the Establishment of Chronic Infection. mBio. 2021;12(2).

39. Jondle CN, Tarakanova VL. T Cell-Intrinsic Interleukin 17 Receptor A Signaling Supports the Establishment of Chronic Murine Gammaherpesvirus 68 Infection. Journal of virology. 2022;96(14):e0063922.

40. Majeed ST, Huss NP, Bradford SM, Jondle CN. B cell-intrinsic interleukin 17 receptor A signaling supports the establishment of chronic murine gammaherpesvirus 68 infection. Journal of virology. 2025;99(12):e0184225.

41. Subramanian S, Tus K, Li QZ, Wang A, Tian XH, Zhou J, et al. A Tlr7 translocation accelerates systemic autoimmunity in murine lupus. Proceedings of the National Academy of Sciences of the United States of America. 2006;103(26):9970–5.

42. Efstathiou S, Ho YM, Hall S, Styles CJ, Scott SD, Gompels UA. Murine herpesvirus 68 is genetically related to the gammaherpesviruses Epstein-Barr virus and herpesvirus saimiri. The Journal of general virology. 1990;71 (Pt 6):1365–72.

43. Virgin HWt, Latreille P, Wamsley P, Hallsworth K, Weck KE, Dal Canto AJ, et al. Complete sequence and genomic analysis of murine gammaherpesvirus 68. J Virol. 1997;71(8):5894–904.

44. Tarakanova VL, Suarez F, Tibbetts SA, Jacoby MA, Weck KE, Hess JL, et al. Murine gammaherpesvirus 68 infection is associated with lymphoproliferative disease and lymphoma in BALB beta2 microglobulin-deficient mice. Journal of virology. 2005;79(23):14668–79.

45. Speck SH, Ganem D. Viral latency and its regulation: lessons from the gamma-herpesviruses. Cell Host Microbe. 2010;8(1):100–15.

46. Usherwood EJ, Roy DJ, Ward K, Surman SL, Dutia BM, Blackman MA, et al. Control of gammaherpesvirus latency by latent antigen-specific CD8(+) T cells. J Exp Med. 2000;192(7):943–52.

47. Tibbetts SA, van Dyk LF, Speck SH, Virgin HWt. Immune control of the number and reactivation phenotype of cells latently infected with a gammaherpesvirus. J Virol. 2002;76(14):7125–32.

48. Usherwood EJ, Ross AJ, Allen DJ, Nash AA. Murine gammaherpesvirus-induced splenomegaly: a critical role for CD4 T cells. The Journal of general virology. 1996;77 (Pt 4):627–30.

49. Collins CM, Boss JM, Speck SH. Identification of infected B-cell populations by using a recombinant murine gammaherpesvirus 68 expressing a fluorescent protein. Journal of virology. 2009;83(13):6484–93.

50. Mboko WP, Olteanu H, Ray A, Xin G, Darrah EJ, Kumar SN, et al. Tumor Suppressor Interferon-Regulatory Factor 1 Counteracts the Germinal Center Reaction Driven by a Cancer-Associated Gammaherpesvirus. Journal of virology. 2015;90(6):2818–29.

51. Collins CM, Speck SH. Expansion of murine gammaherpesvirus latently infected B cells requires T follicular help. PLoS pathogens. 2014;10(5):e1004106.

52. Collins CM, Speck SH. Interleukin 21 signaling in B cells is required for efficient establishment of murine gammaherpesvirus latency. PLoS pathogens. 2015;11(4):e1004831.

53. Sangster MY, Topham DJ, D’Costa S, Cardin RD, Marion TN, Myers LK, et al. Analysis of the virus-specific and nonspecific B cell response to a persistent B-lymphotropic gammaherpesvirus. Journal of immunology (Baltimore, Md : 1950). 2000;164(4):1820–8.

54. Getahun A, Smith MJ, Kogut I, van Dyk LF, Cambier JC. Retention of anergy and inhibition of antibody responses during acute γ herpesvirus 68 infection. J Immunol. 2012;189(6):2965–74.

55. Gauld SB, De Santis JL, Kulinski JM, McGraw JA, Leonardo SM, Ruder EA, et al. Modulation of B-cell tolerance by murine gammaherpesvirus 68 infection: requirement for Orf73 viral gene expression and follicular helper T cells. Immunology. 2013;139(2):197–204.

56. Fleisher GR, Collins M, Fager S. Limitations of available tests for diagnosis of infectious mononucleosis. J Clin Microbiol. 1983;17(4):619–24.

57. Dean LS, Threatt AN, Jones K, Oyewole EO, Pauly M, Wahl M, et al. I don’t know about you, but I’m feeling IL-22. Cytokine Growth Factor Rev. 2024;80:1–11.

58. Melchior K, Gerner RR, Hossain S, Nuccio SP, Moreira CG, Raffatellu M. IL-22-dependent responses and their role during Citrobacter rodentium infection. Infection and immunity. 2024;92(5):e0009924.

59. Alcorn JF. IL-22 Plays a Critical Role in Maintaining Epithelial Integrity During Pulmonary Infection. Frontiers in Immunology. 2020;Volume 11 - 2020.

60. Ivanov S, Renneson J, Fontaine J, Barthelemy A, Paget C, Fernandez EM, et al. Interleukin-22 reduces lung inflammation during influenza A virus infection and protects against secondary bacterial infection. Journal of virology. 2013;87(12):6911–24.

61. Pociask DA, Scheller EV, Mandalapu S, McHugh KJ, Enelow RI, Fattman CL, et al. IL-22 is essential for lung epithelial repair following influenza infection. Am J Pathol. 2013;182(4):1286–96.

62. Jarade A, Garcia Z, Marie S, Demera A, Prinz I, Bousso P, et al. Inflammation triggers ILC3 patrolling of the intestinal barrier. Nature immunology. 2022;23(9):1317–23.

63. Victor AR, Nalin AP, Dong W, McClory S, Wei M, Mao C, et al. IL-18 Drives ILC3 Proliferation and Promotes IL-22 Production via NF-κB. Journal of immunology (Baltimore, Md : 1950). 2017;199(7):2333–42.

64. Perusina Lanfranca M, Lin Y, Fang J, Zou W, Frankel T. Biological and pathological activities of interleukin-22. J Mol Med (Berl). 2016;94(5):523–34.

65. Thorley-Lawson DA. Epstein-Barr virus: exploiting the immune system. Nat Rev Immunol. 2001;1(1):75–82.

66. Oropallo MA, Cerutti A. Germinal center reaction: antigen affinity and presentation explain it all. Trends Immunol. 2014;35(7):287–9.

67. Gauld SB, Santis JL, Kulinski JM, McGraw JA, Leonardo SM, Ruder EA, et al. Modulation of B-cell tolerance by Murine Gammaherpesvirus 68 infection: requirement for Orf73 viral gene expression and follicular helper T cells. Immunology. 2013;139(2):197–204.

68. Collins CM, Speck SH. Tracking murine gammaherpesvirus 68 infection of germinal center B cells in vivo. PloS one. 2012;7(3):e33230-e.

69. Houen G, Trier NH. Epstein-Barr Virus and Systemic Autoimmune Diseases. Front Immunol. 2020;11:587380.

70. Tarakanova VL, Stanitsa E, Leonardo SM, Bigley TM, Gauld SB. Conserved gammaherpesvirus kinase and histone variant H2AX facilitate gammaherpesvirus latency in vivo. Virology. 2010;405(1):50–61.

71. Hu Z, Blackman MA, Kaye KM, Usherwood EJ. Functional heterogeneity in the CD4+ T cell response to murine γ-herpesvirus 68. J Immunol. 2015;194(6):2746–56.

72. Darrah EJ, Jondle CN, Johnson KE, Xin G, Lange PT, Cui W, et al. Conserved Gammaherpesvirus Protein Kinase Selectively Promotes Irrelevant B Cell Responses. J Virol. 2019;93(8).

